# Gluten-Free Beer via Hydrodynamic Cavitation Assisted Brewing of Barley Malts

**DOI:** 10.1101/089482

**Authors:** Lorenzo Albanese, Rosaria Ciriminna, Francesco Meneguzzo, Mario Pagliaro

## Abstract

We provide evidence that novel brewing technology based on controlled hydrodynamic cavitation greatly reduces gluten concentration in wort and finished beer. We advance the hypothesis that the degradation of proline, the most recalcitrant among gluten constituents, leads to gluten concentration reduction in the unfermented as well as in the fermenting wort and later during maturation. These findings are significant as the new cavitation-assisted technology could provide coeliac patients and gluten-intolerant people with gluten-free beer of high quality, offering an alternative to existing methods to lower the gluten concentration, which are detrimental to flavor and taste.

## 1. Introduction

With nearly 200 billion liters per year (Amienyo & Azapagic, 2016), beer is the alcoholic beverage most widely consumed around the world. Both its basic ingredients, *i.e.* water, malt or grains, hops and yeasts, and production methods have barely changed over centuries beyond obvious technological improvements and minor ingredients diversification (Ambrosi, Cardozo, & Tessaro, 2014; Pires & Brányik, 2015). Knowledge of the microbiological processes involved in brewing, on the other hand, has steadily grown over the last decades (Bokulich & Bamforth, 2013).

An open and serious issue in today’s huge beer market is gluten, whose presence in beer arises from barley and wheat malts and grains from which most beers are produced, making beer unsuitable for consumption by coeliac disease patients (Hager, Taylor, Waters, & Arendt, 2014). Production and marketing of “very low gluten content” (concentration < 100 mg/L) or “gluten-free” (< 20 mg/L) beers has been increasingly motivated by the growing awareness about coeliac pathology and milder gluten intolerance.

Contrary to most inflammatory disorders, both genetic precursors and exogenous environmental factors trigger the coeliac disease (Sollid, 2002), which develops in susceptible patients because of their intolerance to ingested wheat gluten or related proteins from rye and barley. In particular, the gluten epitopes that are recognized by the immune system in the human intestine are generally very rich in amino acids and gluten components proline and glutamine residues. Most mammalian peptidases/proteases cannot hydrolyze amide bonds when they are adjacent to proline residues.

Fermentation, usually lasting several days, is the most important brewing step for the gluten reduction in traditional beers. It begins with the pitching of yeasts in the quickly cooled and aerated wort, usually from the strain *Saccharomyces cerevisiae* (Albanese, Ciriminna, Meneguzzo, & Pagliaro, 2015), whit initial yeast concentration generally between 15 and 20 million cells per mL. In today’s most used cylinder-conical, closed and thermostatically controlled fermenters, early assimilation of fermentable sugars, amino acids, minerals and other nutrients occurs along with metabolic production of ethanol, CO_2_, higher alcohols, esters and other substances (Bokulich & Bamforth, 2013). Such fermentation products, while often toxic to yeasts at high concentration, are desirable for the quality of the certain beer styles as well contribute to their aromatic bouquet (Pires & Brányik, 2015). Among them, several esters providing characteristic beer aromas are synthesized via a biochemical pathway involving ethanol and higher alcohols (Landaud, Latrille, & Corrieu, 2001).

Few attempts aimed at modeling the complex fermentation processes, despite encouraging results (Albanese et al., 2015), are intrinsically limited by the extreme dependency on few factors hardly generalizable such as yeast strain, aeration and CO_2_ concentration occurring in the full beer brewing environment (Brown & Hammond, 2003). In general, effective fermentation is particularly relevant for the degradation of amino acids accumulated in the fermenting wort, supplying nearly all the nitrogen needed by the yeasts’ cellular biosynthesis in the form of free amino-nitrogen (FAN), as well as affecting bitterness, flavor and foam stability (Choi, Ahn, Kim, Han, & Kim, 2015).

The most important FAN is glutamine – an amino acid and gluten component – along with other amino acids belonging to the so called “Group A” which undergo the fastest assimilation by yeast cells at a rate depending on the specific yeast strain (Pires & Brányik, 2015). Once Group A amino acids are assimilated, other ones belonging to Groups B and C are more gradually and slowly assimilated until nitrogen-deprived residuals from original amino acids are turned into higher (fusel) alcohols and esters, strongly impacting beers’ flavor. Only one amino acid belongs to Group D, namely proline – another basic gluten component – whose assimilation by yeast cells was deemed negligible until few years ago (Lekkas, Stewart, Hill, Taidi, & Hodgson, 2005).

Most if not all production of gluten-free beers foresees the use of at least a fraction of malts derived from cereals and pseudo-cereals not containing gluten or its precursors, such as sorghum, buckwheat, quinoa, amaranth (Meo et al., 2011; Wijngaard & Arendt, 2006), maize and oat (Yeo & Liu, 2014). However, the respective brewing techniques for cereals different from barley are not yet well established. Recently, encouraging results were achieved with rice malt brewing in a pilot plant, affording an alcoholic product with a sensory profile similar to a barley malt beer in aroma, taste and mouthfeel, though flatter and featuring rapidly collapsing foam (Mayer et al., 2016).

In alternative, complex and costly filtration and enzymatic techniques are used, to condition the malts in order to boost processes leading to the precipitation of proteins, during mashing, fermentation and stabilization (Dostálek, Hochel, Méndez, Hernando, & Gabrovská, 2006; Hager et al., 2014). Another alternative technique consists in the use of silica gel (SG) in the fermentation stage of the brewing process to selectively remove proteins without practically affecting both valuable yeast nutrients such as free amino-nitrogen (FAN) and foam-causing proteins, leading to gluten reduction in the subsequent stabilization stage (Benítez, Acquisgrana, Peruchena, Sosa, & Lozano, 2016). Although silica is generally recognized as a safe (GRAS) food additive both in US and Europe, its use adds to cost and process complexity. Beyond uncertainties, complexity and costs, the resulting finished beers most often have different aroma and flavor when compared to conventional beers.

Now, we show the first evidence of the potential for brewing of conventional barley malt assisted by controlled hydrodynamic cavitation (CHC) to reduce the gluten concentration in beer by means of suitable cavitation regimes and operational parameters, *i.e.* by purely electro-mechanical means, without either changing ingredients or using additives as well as any other chemical or biochemical solution. The CHC process, including the oxidation processes triggered by hydrodynamic cavitation, is the one we recently developed and tested with a real-scale experimental demonstration device, finding significant advantages including *i*) dramatic reduction of saccharification temperature; *ii*) increased and accelerated peak starch extraction; *iii*) significant reduction of operational time eliminating traditional stages such as dry milling and boiling; *iv*) relevant energy saving; *v*) shorter cleaning time; *vi*) volumetric heating which prevents caramelization; and *vii*) overall simplification of both structural setup and operational management of brewing processes (Albanese, Ciriminna, Meneguzzo, & Pagliaro, 2016).

## 2. Materials and methods

### 2.1. Brewing units

Among controlled hydrodynamic cavitation devices in which hydrocavitation regimes can be controlled and tuned so as to avoid damage to the equipment, Venturi-type stationary reactors are the most appealing candidates for industrial-scale applications due to their cheapness as well as intrinsic ease of construction, scale-up, and reduced risk of obstruction from solid particles and other viscous substances such as those found in brewing (Albanese et al., 2015). Hence, a dedicated equipment was built from known or commonly available commercial components, in order to investigate the effects of hydrodynamic cavitation processes upon gluten concentration.

Figure 1 shows the device used for the experimental tests, including a closed hydraulic loop with total volume capacity around 230 L, powered by a centrifugal pump (Lowara, Vicenza, Italy, model ESHE 50-160/75 with 7.5 kW nominal mechanical power) with open impeller 0.174 m in diameter. Rotation speed was set around 2900 rpm.

**Figure 1.**
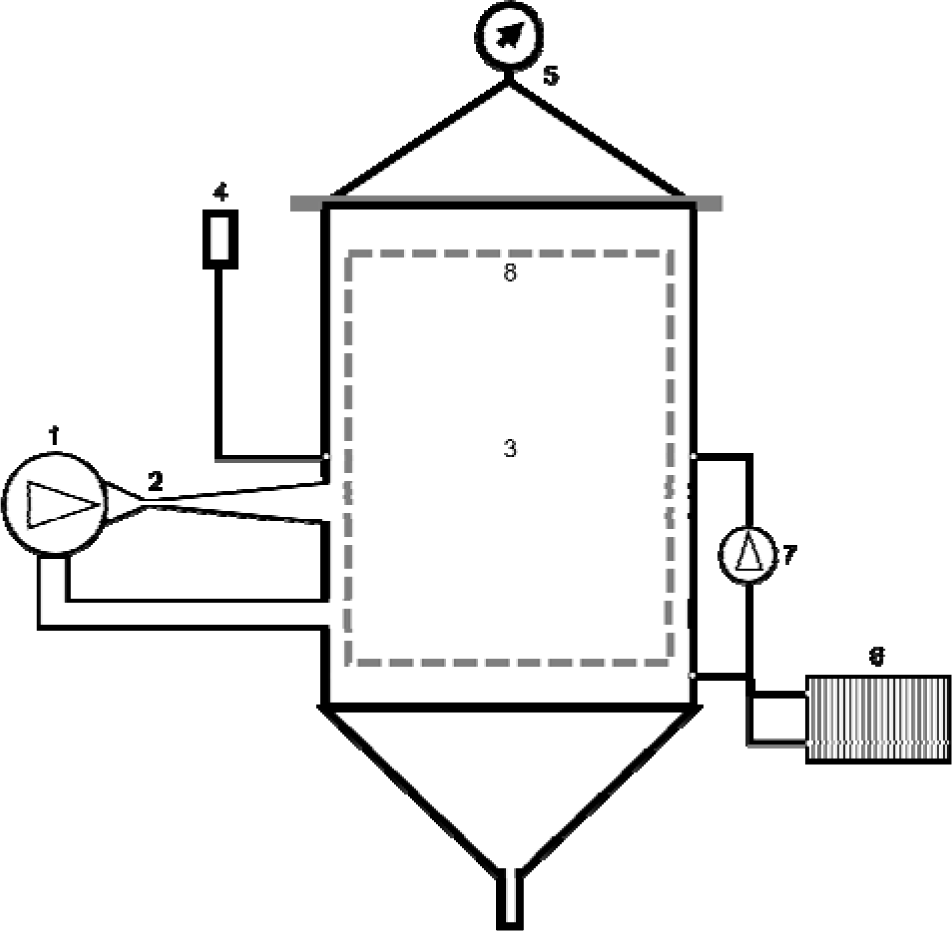
Simplified scheme of the experimental CHC-based installation. 1 – centrifugal pump, 2 – CHC reactor, 3 – main vessel, 4 – pressure release valve, 5 – cover and manometer, 6 – heat exchanger, 7 – circulation pump, 8 – malts caging vessel. Other components are commonly used in state-of-the-art hydraulic constructions.

Any surface in contact with the wort was crafted in food-quality stainless steel (AISI 304), with 2 mm minimum thickness. The circulating liquid (wort) can be exposed to the atmospheric pressure or to a given average pressure limited by a tunable pressure release valve. Such valve was preferred over an expansion tank in order to avoid wort contamination by substances accumulated in the tank during successive tests, while performing the same task, *i.e.* tuning the cavitation intensity through the *P_0_* term in the Bernoulli’s equation, shown in its simplest form by Equation 1:

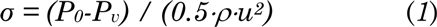

where *σ* is the cavitation number (also indicated as CN), *P_0_* (Nm^-2^) is the average pressure downstream of a cavitation reactor such as a Venturi tube or an orifice plate in which the cavitation bubbles collapse, *P_v_* (Nm^-2^) is the liquid vapor pressure (a function of the average temperature for any given liquid), *ρ*(kgm^-3^) is the liquid density, and *u* (ms^-1^) is the flow velocity through the nozzle of the cavitation reactor.

Different CHC regimes are practically achieved according to the values assumed by the cavitation number in one of the three intervals, corresponding to broad cavitational regimes, identified by Gogate (Bagal & Gogate, 2014; Gogate, 2002), ideally, without impurities and dissolved gases, cavities would be generated for values of *σ* < 1. For *σ* < 0.1 values, the cavities are no longer able to collapse and the CHC regime turns to chocked cavitation or supercavitation. For *σ* > 1, lesser and lesser cavities are generated, while their collapse becomes ever more violent. For the scope of this study, only the developed cavitation with 0.1<*σ*<1 will be considered.

Volumetric liquid heating occurs during circulation due to the conversion of impeller’s mechanical energy into thermal energy, particularly downstream the cavitation reactor nested into the hydraulic loop, due to the vigorous internal friction associated with the cavitation process. As a closed hydraulic circuit, no change of potential energy is involved, therefore all the mechanical energy turns into heat.

A Venturi tube, whose geometry was described in a previous study (Albanese et al., 2015), was used as the cavitation reactor and preferred over an orifice plate since orifices are quickly obstructed by the circulating solid particles. Moreover, a smaller circulation pump (Rover pompe, Padova, Italy, model NOVAX 20 B, power 340 W, working temperature up to 95°C, capacity up to 28 L min^-1^), drives a secondary recirculation loop through an ordinary plate heat exchanger (20 stainless steel plates, each with a 0.043 m^2^ surface area), allowing for isothermal stages when required in the course of the brewing process, depending on specific brewing recipes. The latter pump was used as well both to cool the wort after brewing and to convey it to the fermenters. Onboard sensors included a manometer and few digital thermometers (not shown), hydraulic pressure and temperature being the main physical parameters monitored and actually used to manage the brewing processes.

The installation was designed to perform the mashing and hopping stages of brewing, while fermentation was generally performed in common 200 L stainless steel cylinder-conical fermenters after receiving the wort from the main unit. Though, a few tests were performed without wort removal, using the installation shown in Figure 1 as the fermenter.

All but one of the tests designed to study the CHC effects upon the gluten concentration were run in brew in the bag (BIAB) mode, thus using the malts caging vessel in Figure 1. In one test, the vessel was absent and malts were allowed circulating. Water, before being conveyed to the brewing unit, is passed through a mechanical filter made up of a 20 μm polypropylene wire to remove solid particles down to 50 μm in size. An active carbon filter reduces chlorine concentration, attenuates odors and flavors, and removes other impurities down to 70 μm in size. The pH is usually lowered from about 7 to about 5.5 by adding 80 wt% lactic acid (70-80 mL). For the purpose of comparison, traditional brewing was performed by means of a Braumeister (Ofterdingen, Germany) model 50 L brewer, equipped with a cooling serpentine (model Speidel stainless steel wort chiller for 50-liter Braumeister) and fully automatic brewing control (temperature, time and recirculation pumps). Finally, after fermentation, bottling was performed via an ordinary depression pump (Tenco, Genova, Italy, model Enolmatic, with capacity around 200 bottles per hour).

### 2.2. Analytical instruments and methods

Along with thermometer and manometer sensors onboard the main production unit, few specialized off-line instruments were used to measure the chemical and physiological properties of wort and beer. The acidity was measured by means of pH-meter (Hanna Instruments, Padova, Italy, model HI 99151) with automatic pH calibration and temperature compensation. The sugar concentration in the wort during mashing and before fermentation was measured in Brix percentage degrees by means of a refractometer (Hanna Instruments, Padova, Italy, model HI 96811, scaled from 0% to 50% Brix, resolution 0.1%, precision ±0.2% in the 0-80°C temperature range, and automatic temperature compensation in the 0-40°C range). Brix readings were then converted to starch extraction efficiency (Bohačenko, Chmelík, & Psota, 2006).

The specific sugar content of any used malt was multiplied by its mass in the wort to produce the overall sugar content, which is in turn directly related to the peak theoretical apparent specific gravity (ratio of the weight of a volume of the substance to the weight of an equal volume of water) at 20°C according to widely available data tables. The real specific gravity was computed from the Brix reading (again at 20°C), with starch extraction efficiency represented by the ratio of the latter to the peak theoretical apparent specific gravity.

Physico-chemical and physiological parameters of fermenting wort and finished beer were measured by means of a 6-channel photometric device (CDR, Firenze, Italy, model BeerLab Touch). Relevant to this study were fermentable sugars (0.1 to 150 g/L of maltose, resolution 0.01 g/L), free amino-nitrogen, or FAN (30 to 300 mg/L, resolution 1 mg/L), alcohol content (0-10% in volume, resolution 0.1%). All reagents were of analytical grade. The gluten concentration measurement method was RIDASCREEN Gliadin competitive, *i.e.* the official standard method for gluten determination according to the Codex Alimentarius (Hager et al., 2014; Rallabhandi, Sharma, Pereira, & Williams, 2015). The results were in units mg/L with an upper limit at 270 mg/L and the measurement uncertainties, as declared by the accredited laboratory in charge of the analyses, were equal to 6.7% for results above 150 mg/L, as low as 2.4% below such threshold.

The microbiological measurement, *i.e.* the counting of the living yeast cells, was performed according to the method explained in the Appendix to our previous study (Albanese et al., 2015). Electricity consumption and power absorption by the centrifugal pump were measured by means of a commercial digital power meter. Once the water flow was set, its speed through the Venturi’s nozzle was computed from straightforward division by its section, hence the cavitation number as per Equation (1). Water density (10^3^ kg/m^3^) was used throughout all calculations, which may lead to a small underestimation of the CN (malt and starch are less dense than water).

### 2.3. Brewing ingredients

Pilsner or Pale were used as the base barley malts in all experiments, along with smaller fractions of Cara Pils, Cara Hell and Weizen. Among the hops, different combinations of pelletized German Perle, Saaz and German Hersbrucker were used. In the course of few tests, candied brown sugar was added to the wort before fermentation, while regular white sugar was added to the fermented wort before bottling and maturation, aimed at carbonation. Finally, fermentation was activated by means of the dry yeast strain Safale US-05, requiring temperature between 15°C and 24°C and maximum alcohol content 9.2%, used in any test in the identical proportion of 67 g per 100 L. As a relevant exception, test B2 was carried out with 47 g of yeast, namely in the proportion of 94 g per 100 L (40% greater than in other tests).

### 2.4. Production tests

Table 1 summarizes few basic features of the brewing tests reported in this study. No simple sugar was added during the mashing stage in any test. For three of such tests (C8, C9 and C10) sharing the same ingredients, the fermentation step was performed in the device. Finally, two tests (B1 and B2) were performed with a traditional equipment.

**Table 1.** Beer production tests, ingredients and conditions.

| Test ID | Brewing unit<sup>a</sup> | Net volume (L) | Malt | Cavitating malts | Hops<sup>b</sup> | Added sugars<sup>d</sup> | Fermentation<sup>e</sup> |
| --- | --- | --- | --- | --- | --- | --- | --- |
| CO1 | 1(A) | 186 | Pilsner 25 kg<br>Cara Pils 1.6 kg<br>Cara Hell 2.6 kg<br>Weizen 2 kg | Yes | Perle 0.1 kg<br>Hers<sup>c</sup> 0.3 kg<br>Saaz 0.2 kg | W 0.96 kg (bot) | Standard |
| C2 | 1(B) | 170 | Pilsner 25 kg<br>Cara Pils 1.6 kg<br>Cara Hell 2.6 kg<br>Weizen 2 kg | No | Perle 0.1 kg<br>Hers 0.4 kg<br>Saaz 0.1 kg | W 0.96 kg (bot) | Standard |
| C5 | 1(B) | 170 | Pilsner 25 kg<br>Cara Pils 3.6 kg<br>Cara Hell 2.6 kg | No | Perle 0.1 kg<br>Hers 0.3 kg<br>Saaz 0.2 kg | W 10 kg (fer)<br>W 0.96 kg (bot) | Standard |
| C6 | 1(B) | 170 | Pilsner 25 kg<br>Cara Pils 3.6 kg<br>Cara Hell 2.6 kg | No | Perle 0.3 kg<br>Hers 0.4 kg<br>Saaz 0.2 kg | W 8 kg (fer)<br>W 0.96 kg (bot) | Standard |
| B1 | B-50 | 50 | Pilsner 6.25 kg<br>Cara Pils 0.9 kg<br>Cara Hell 0.65 kg | No | Perle 0.025 kg<br>Hers 0.075 kg<br>Saaz 0.05 kg | W 0.042 kg (bot) | Standard |
| C7 | 1(B) | 170 | Pilsner 28.5 kg<br>Cara Pils 2.5 kg | No | Perle 0.6 kg<br>Saaz 0.5 kg | W 0.84 kg (bot) | Standard |
| B2 | B-50 | 50 | Pilsner 9.2 kg<br>Cara Pils 0.81 kg | No | Perle 0.194 kg<br>Saaz 0.162 kg | W 0.07 kg (bot) | Standard |
| C8 | 1(B) | 170 | Pale 26 kg<br>Cara Pils 3 kg | No | Perle 0.2 kg<br>Hers 0.1 kg | B 1.0 kg (fer)<br>W 0.96 kg (bot) | Device |
| C9 | 1(B) | 170 | Pale 26 kg<br>Cara Pils 3 kg | No | Perle 0.2 kg<br>Hers 0.1 kg | B 1.0 kg (fer)<br>W 0.96 kg (bot) | Device |
| C10 | 1(B) | 170 | Pale 26 kg<br>Cara Pils 3 kg | No | Perle 0.2 kg<br>Hers 0.1 kg | B 1.0 kg (fer)<br>W 0.96 kg (bot) | Device |

a) With reference to Figure 1: 1(A) = circulating malts (no caging vessel); 1(B) = with caging vessel (BIAB mode). B-50 = Braumeister model 50-liters.
b) Hops cavitating in any test.
c) Hers = Hallertau Hersbrucker hop.
d) W = simple white sugar; B = candied brown sugar; bot = before bottling; fer = before fermentation.
e) Standard = fermentation in cylinder-conical vessel 200 l. Device = fermentation performed into the experimental device.

## 3. Results and Discussion

The tests carried out by means of CHC-assisted brewing led to significant reduction of gluten final concentration in the produced beers that, to the best of our knowledge, represents a novelty. Figure 2 shows the gluten concentration for all the performed tests including the fermentation stage, measured at different times starting at the beginning of fermentation, and during maturation (generally performed in bottles). Although barley malts were used practically in the same proportion to the respective volumes (Table 1), large differences are striking, with tests C6, B2, C8, C9, C10, and partially test C7, showing far less final gluten concentration.

**Figure 2.**
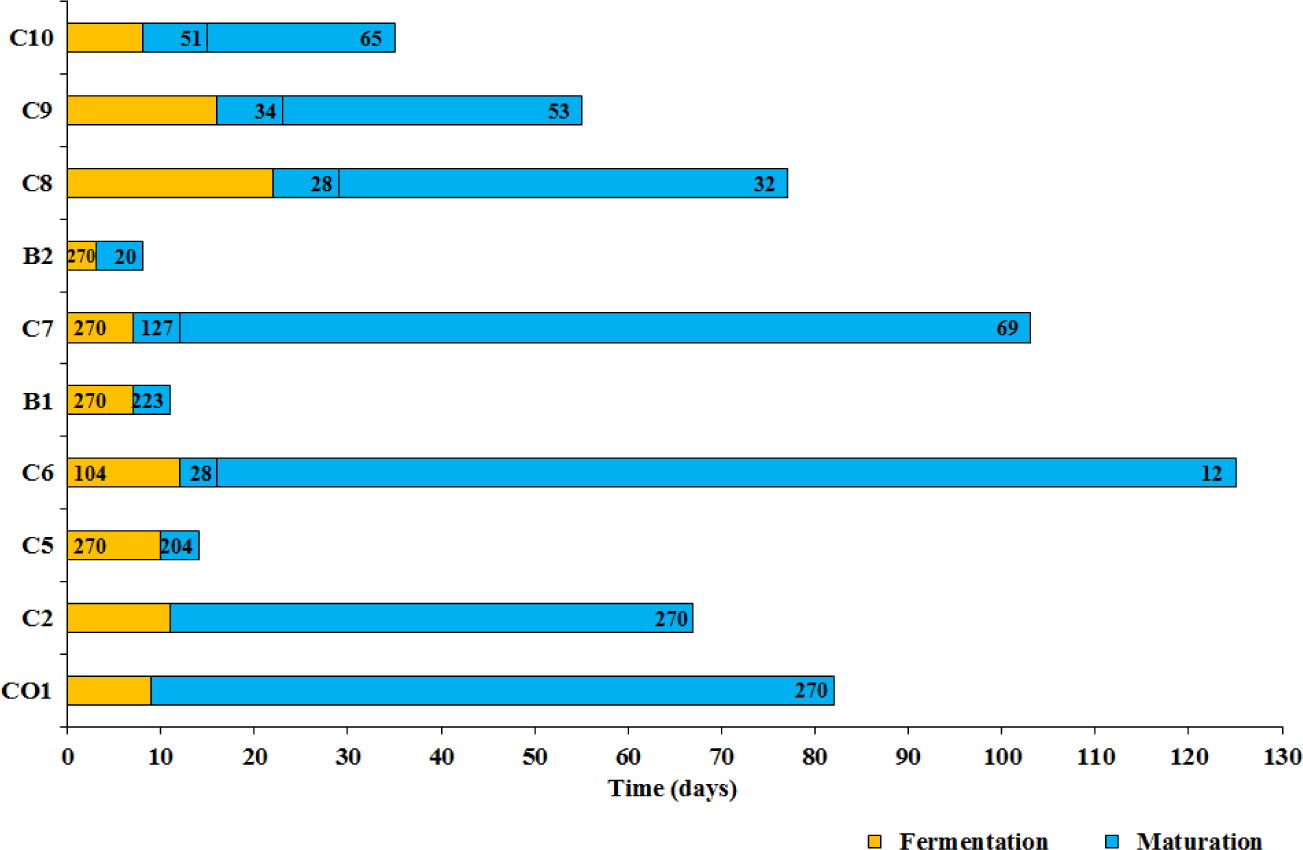
Gluten concentration at different times after the brewing production processes. The value of 270 mg/L should be considered as the lower limit. Uncertainties not shown.

In tests C8 and C10, differently from all others, CHC processes were activated also after yeasts pitching and the subsequent fermentation occurred in the installation shown in Figure 1, as well as in test C9 for the purpose of comparison with tests C8 and C10. Moreover, in these latter tests, Pale malt replaced Pilsner (see Section 2.3).

Tests C5 and C6 were carried out with exactly the same malts, while the net process times before fermentation were different resulting in electricity consumptions equal to 84 kWh and 108 kWh, respectively, far greater than in tests CO1 (60 kWh) and C2 (28 kWh, terminated at the temperature of 78°C). Starch extraction efficiencies in tests C5 and C6 were nearly identical too, *i.e.* 57% and 56%, respectively (see Figure 4 in Albanese et al., 2016), as well as the fermentation times (10 and 12 days, respectively), while yeast strains and their respective quantities were identical. Nevertheless, test C6 showed a gluten concentration far lower than test C5 at any time: 104±2.5 mg/L vs >270±18 mg/L at the beginning of fermentation, and 28 mg/L (*i.e.* just above the gluten-free threshold) *vs.* 204 mg/L after approximately the same time after test (14 and 16 days, respectively).

Eventually, 125 days after brewing (excluding fermentation), beer produced by means of test C6 fell well below the gluten-free threshold, down at a mere 12±0.3 mg/L concentration.

The brewing processes used in tests C5 and C6, in terms of temperature and cavitation number (CN) against consumed energy is summarized in Figure 3(a), while Figure 3(b) shows the gluten concentration values in the course of the brewing processes carried out in the same tests before fermentation.

**Figure 3.**
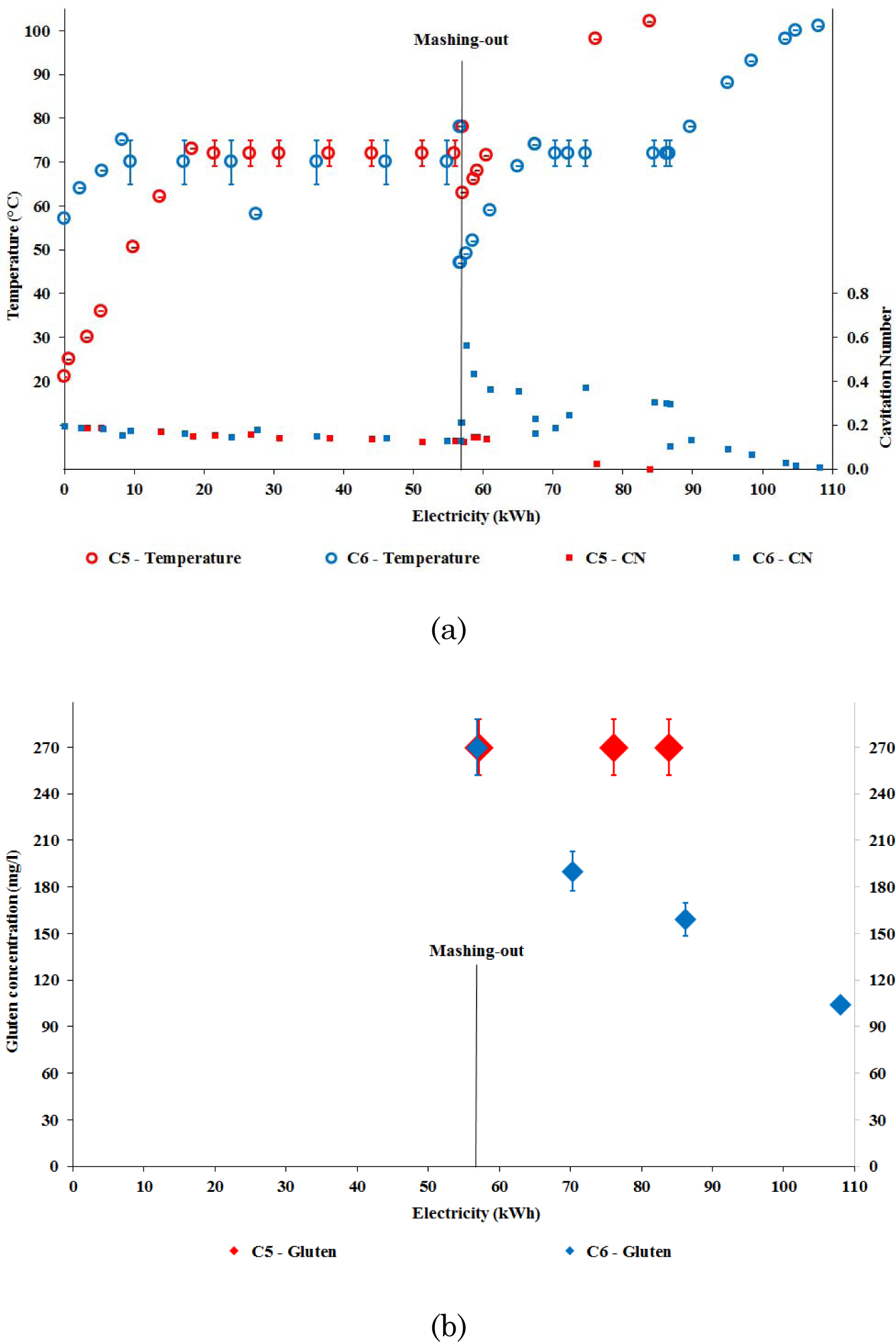
Tests C5 and C6: temperature with respective uncertainties and cavitation number (a), and gluten concentration with respective uncertainties (b), both reproduced as a function of consumed electricity.

The differences in the initial temperature of each process before mashing-out (21°C in test C5 and 57°C in test C6) did not lead to any effect on gluten concentration at the time of mashing-out (both >270 mg/L).

After restart following mashing-out, an additional hydraulic pressure was readily imposed in C6 process, oscillating between 0.5 atm and 2 atm and averaging 1.5 atm, with approximately 30 kWh energy consumed during the overpressure stage, keeping the temperature at around 72°C (almost 20 kWh of consumed electricity). This resulted in far greater CN values for test C6, more than double the values at similar temperatures before the activation of the additional pressure, hinting to a more violent hydraulic cavitation regime. Rather surprisingly, Figure 3(b) shows that the gluten concentration for test C6 fell far below the values observed for test C5 at the same energy consumption (always >270±18 mg/L for test C5, while in test C6 it decreased to about 160±11 mg/L at the energy consumption of about 84 kWh). Eventually, the gluten concentration in test C6 was only 104±2.5 mg/L before fermentation, with overall energy consumption amounting to 108 kWh.

In tests CO1 and C2, with type and quantities of malts very similar to tests C5 and C6, no additional pressure was applied and the net process times were significantly smaller than in tests C5 and C6. Figure 2 shows that the gluten concentrations in the respective beers were > 270±18 mg/L after 60 to 80 days since the beginning of fermentation.

Test C7 involved the use of a marginally different dosage of malts, but the same overall quantity as in tests C5 and C6, as shown in Table 1. Moreover, since the mashing efficiency was far greater in test C7 (71%) than in tests C5 and C6 (Albanese et al., 2016), one would expect that – all else being equal – the gluten concentration in the finished beer resulting from test C7 will be higher. No additional hydraulic pressure was ever applied, so that cavitation numbers were very close to those assessed during brewing in test C5. Actually, the gluten concentration in test C7 was always >270±18 mg/L during brewing before fermentation, similar to test C5. Moreover, the fermentation time in test C7 was equal to 7 days, shorter than in both tests C5 (10 days) and C6 (12 days). One might therefore wonder why the gluten concentration in the beer from test C7, measured during maturation, was far lower than in test C5 just 5 days (C7) and 4 days (C5) after the end of fermentation, *i.e.* 127±3 mg/L against 204±14 mg/L. Eventually, 103 days after brewing, beer produced by means of test C7 fell well below the very low gluten content threshold, at just 69±2 mg/L.

Figure 4 shows the brewing processes used in test C7, again in terms of temperature and cavitation number (CN), against consumed energy. While the overall process time before fermentation was similar to tests C5 and C6, the starch extraction is now far more efficient, resulting in mashing-out performed in correspondence of an energy consumption equal to 27 kWh, against about 57 kWh for tests C5 and C6.

This translates into less than half time elapsed from process beginning. Therefore, a much larger fraction of the overall process time occurs after mashing-out, *i.e.* with all the starch and the gluten content being available in the wort and undergoing the hydraulic cavitation processes.

The main hypothesis we advance to explain the above results involves the generation of molecular oxygen by means of the water splitting reactions triggered by hydrodynamic cavitation (Ciriminna, Albanese, Meneguzzo, & Pagliaro, 2016b), represented by Equations 2-7:

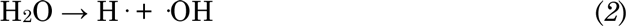

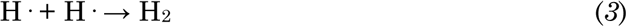

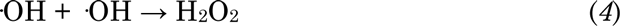

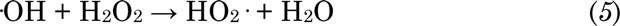

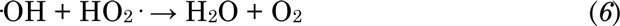

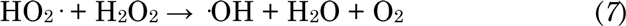

Contrary to acoustic cavitation in which the formation, growth, and implosive collapse of cavitation bubbles is caused by the propagation of ultrasonic waves (Mason & Peters, 2002), in hydrodynamic cavitation the same phenomena are produced by the motion of fluid, with all the liquid being forced through an aptly developed cavitation reactor (Venturi tube, orifice plate, rotor-stator and others). Constrictions and nozzles, resulting in acceleration and local depressurization, alter the flow geometry. If the pressure falls below the boiling point, water vaporizes and vapor bubbles are generated, whose collapse in *vena contracta* downstream the nozzle generates local hot spots with extreme temperature (>5000 K), pressure (>200 atm) and hydraulic jets (>150 m/s), as well as splitting of the water molecules into H_2_, O_2_, and hydroxyl radicals (·OH) (Yasui, Tuziuti, Sivakumar, & Iida, 2004).

Equation (6) and equation (7) are especially important in this context. The O_2_ generation occurring during cavitation may activate the enzyme oxidase (Schnitzenbaumer & Arendt, 2014) in the course of mashing, whereas its persistence in the fermenting wort does the same later with yeast oxidase (Procopio et al., 2013). Furthermore, the proline molecules might be degraded or at least partially destroyed by the extreme thermo-mechanical events triggered under violent cavitation regimes such as those activated in test C6, as well as by milder cavitation regimes, *i.e.* occurring at atmospheric pressure, provided that such regimes are enabled for a sufficient time interval, such as in test C7.

Based on the performed tests and the above hypotheses, CN > 0.3 during a sufficient time seems to be required for the destruction of proline molecules, whereas data in Figure 3(a) suggest an upper limit to such time sufficient to obtain gluten-free beer, in terms of specific consumed energy, of about 0.17 kWh/L (resulting from about 30 kWh divided by 170 liters). While the identification of a lower limit to achieve gluten-free beer will require further tests, it is conceivable that such limit could be significantly lower, as Figure 2 shows that the later gluten concentration in test C6, 125 days after brewing (excluding fermentation), was just 12±0.3 mg/L.

Concerning milder cavitation regimes (test C7), data in Figure 4 shows that a similar upper limit to their activation time, in terms of specific consumed energy and sufficient to obtain very low gluten content beer, of about 0.29 kWh/L (resulting from about 50 kWh divided by 170 liters). Again, the identification of a lower limit to achieve very low gluten content beer will require further tests, though it is conceivable that such limit could be significantly lower, as Figure 2 shows that the later gluten concentration in test C7, 103 days after brewing (excluding fermentation), was just 69±2 mg/L.

**Figure 4.**
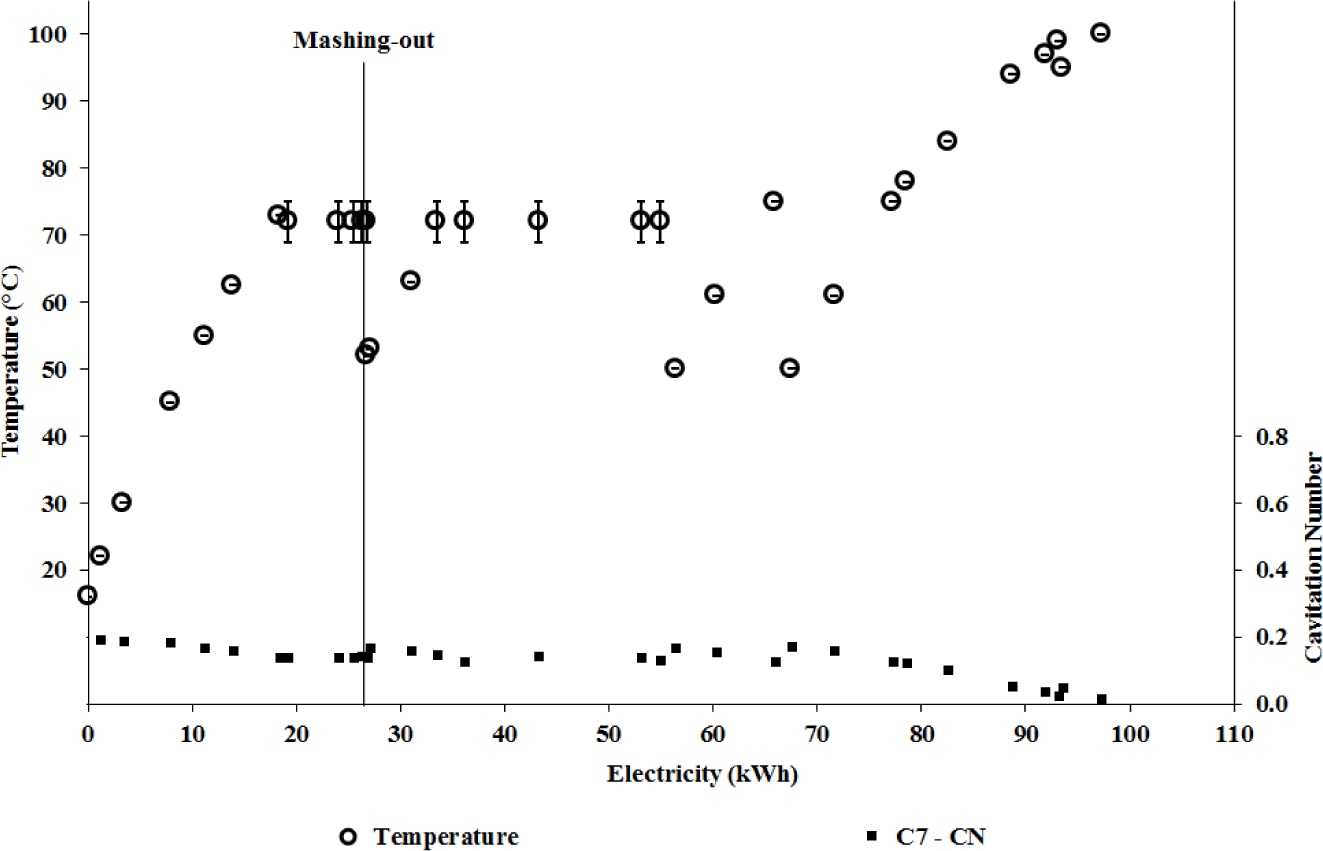
Same as Figure 3(a), for test C7.

The observed sustained decline of gluten concentration during maturation in tests C6 and C7 shown in Figure 2, along with the consideration that no filtration was performed and most food and nutrients for yeasts remained available, might imply that the yeasts remain active and assimilating proline during maturation and that molecular oxygen persists in the fermenting wort.

Whatever the underlying mechanisms triggered by the hydrodynamic cavitation, the effect in terms of reduced gluten concentration is similar to complex chemical techniques (Dostálek et al., 2006). Gluten data for tests C5, C6 and C7 after brewing, although already shown in Figure 2, are better appreciated through Figure 5, showing the concentration decay in terms of absolute values as well as percentage of initial values, with test C6 clearly outperforming all the others.

**Figure 5.**
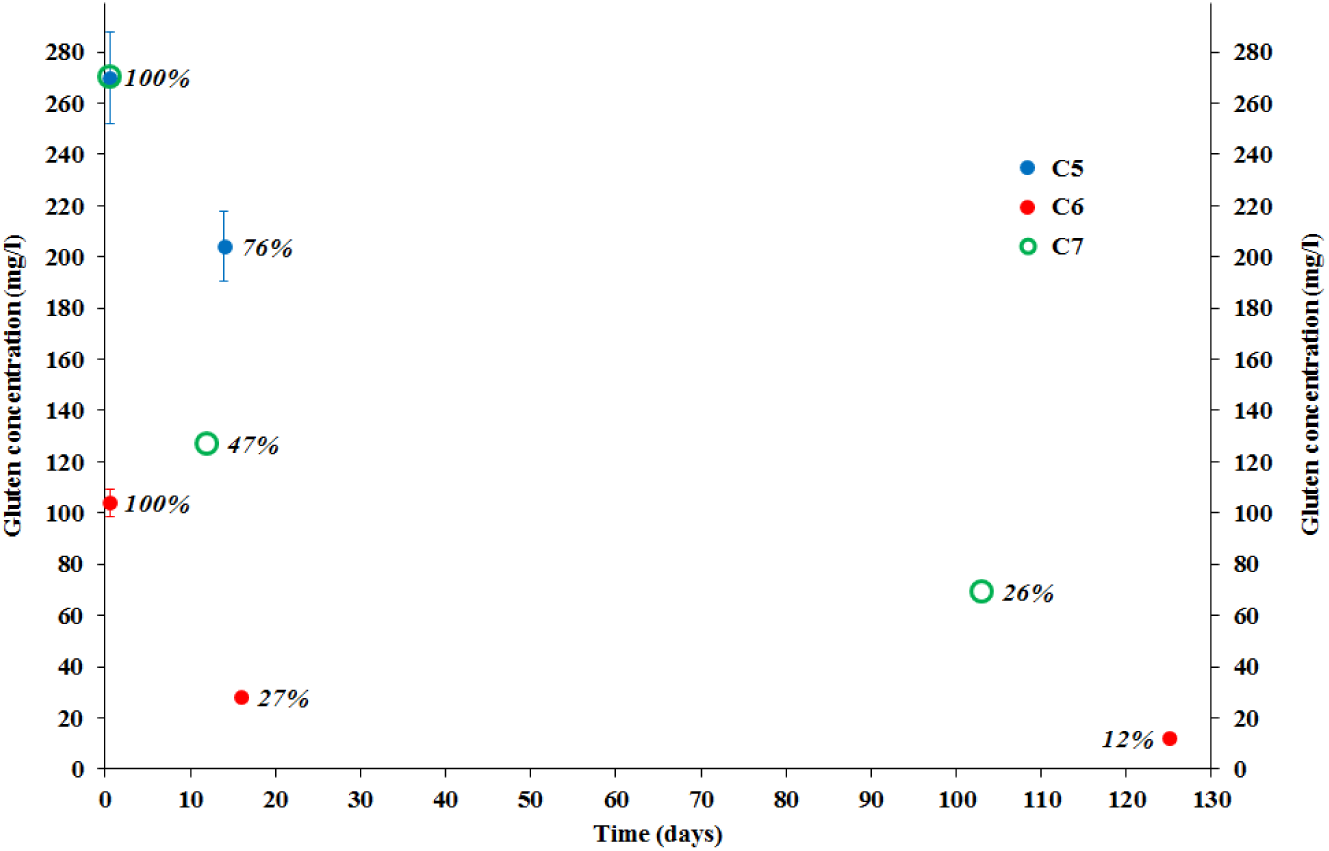
Tests C5, C6 and C7: decay of gluten concentration as a function of time after the respective brewing production processes.

The three-data gluten concentration series concerning tests C6 and C7 are best fitted by a power decay function of time, with more than 99.5% of variance explained. In the event that such relationship could be proved to hold in general, it would provide predictive capability about the eventual achievement of either the very low gluten content (100 mg/L), or gluten-free (20 mg/L) thresholds, starting from given initial conditions, brewing recipe and operational parameters.

As shown in Table 1, tests B1 and B2 were carried out by means of the conventional B-50 installation, with test B2 involving supply of a quantity of barley malts about 28% greater than B1, and showing higher starch extraction efficiency too, *i.e.* 71% vs 60% (Albanese et al., 2016). Therefore, all else being equal, one might expect a greater gluten concentration from test B2. On the contrary, beer from such test showed a gluten concentration as little as 20±0.5 mg/L after 5 days of maturation following a mere 3-days fermentation, against 223±15 mg/L for test B1 after 7 days of fermentation followed by 4 days of maturation.

The achievement of the above-mentioned very low gluten concentration in test B2 was most likely a consequence of the far greater yeast concentration supplied to the fermentation stage when compared to test B1. The feasibility of such method for gluten reduction would be anyway undermined by the significant harm to the beer taste, flavor and aroma produced by the greater yeast supply, as was indeed subjectively observed for the beer resulting from test B2, let alone the greatly increased process time and energy consumption.

Turning to tests C8, C9 and C10, Figure 2 shows that the respective gluten concentrations are relatively low, which is especially relevant for test C9 when no additional cavitation process was activated after yeast pitching, thereby confirming the expectations based upon the use of Pale malt in place of Pilsner one (see Section 2.3). Recalling that those tests were carried out with exactly the same ingredients, Figure 6 shows that the brewing processes up to mashing-out were practically identical in terms of energy consumption in both tests C8 and C9 and (4 kWh less consumed energy) in test C10. Up to the yeast pitching, the consumed energy in those tests was far lower than in tests C5, C6 and C7. Moreover, all three tests were carried out at atmospheric pressure, producing the same cavitation numbers at the corresponding temperatures.

**Figure 6.**
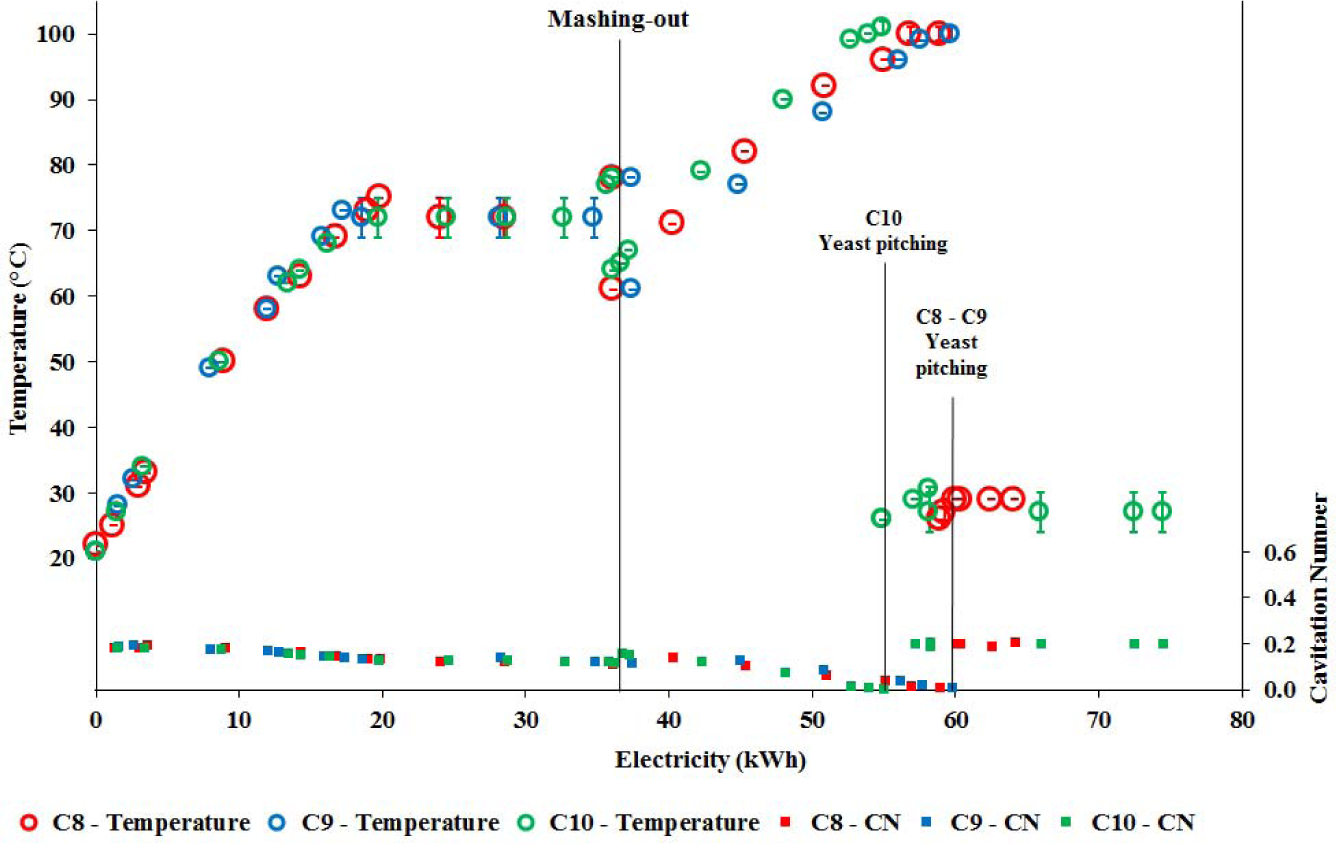
Same as Figure 3(a), for tests C8, C9 and C10.

After yeast pitching, CHC was activated in both tests C8 and C10, with energy consumption around 5 kWh and 19.5 kWh, respectively, while no CHC was applied to test C9. The main motivation for such further tests was the need to check the sensitivity to the replacement of the Pilsner with the Pale malt as the main mashing ingredient, as well as the analysis of any possible advantage brought by the cavitation processes after yeast pitching, as claimed in a previous study (Safonova, Potapov, & Vagaytseva, 2015).

Figure 2 shows that the overall gluten concentration in test C9 was indeed significantly lower – below the very low gluten content threshold – than in tests C5 and C7, despite the smaller energy consumption during brewing in test C9.

However, Figure 7 hints to a definite relationship between residence time in the open vessel (indicated as “fermentation”, even if strictly speaking fermentation could have stopped well before the time of bottling, at least for tests C8 and C9) and gluten concentration measured 7 days after bottling.

**Figure 7.**
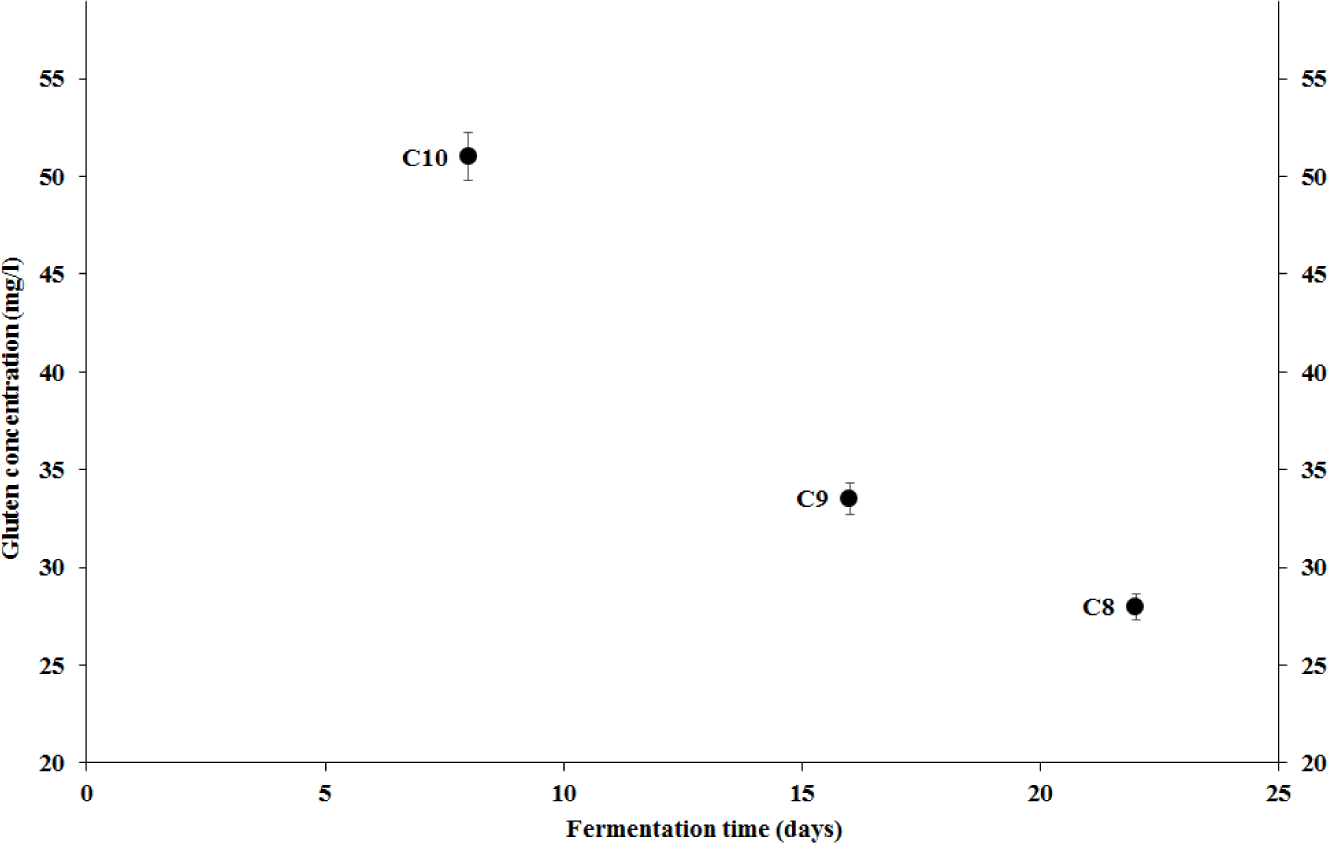
Tests C8, C9 and C10: gluten concentration measured 7 days after bottling, as a function of the length of the fermentation period.

Regardless of any CHC treatment after yeast pitching, the gluten concentration decreases with fermentation time according to a power decay function of time, with more than 99.9% variance explained, despite its significance cannot be established based on only three data points. Such relationship seems to agree with the consideration expressed above as well as in Section 1 about the sensitivity of the proline assimilation rate to the availability of molecular oxygen, which increases with the residence time in an open vessel (Procopio et al., 2013).

Another rather surprising feature, shown again by Figure 2, is the absence of any decay in gluten concentration during maturation: on the contrary, an increase with time was observed, largest in test C9 (from 34±1 mg/L to 53±1 mg/L during 32 days of maturation in bottles), and insignificant in test C8. We ascribe such growth to the possible release of FAN during maturation from inactivated or dead yeast cells, along with the additional hypothesis that a significant fraction of such released FAN is non-degraded glutamine or – more likely – proline, *i.e.* gluten constituents. The hypothesis is supported by similar recent findings in which a sample of wort beer after yeast pitching treated in an ultrasonic bath of given frequency and variable power (Choi et al., 2015) led to acceleration of FAN utilization by yeasts (in its turn improving the beer’s organoleptic and physiological properties).

While it remains unclear why the same phenomenon is not shown in any other test, given that yeasts did not undergo any cavitation process in test C9, the larger increase of gluten concentration in test C9 in comparison with tests C8 and C10 could hint to a relevant role of yeast cells “activation” by CHC processes.

More in detail, Figure 8 shows that, during the residence in the open vessel (fermentation stage), an inverse relationship seems to hold between the tendencies of the concentration of yeast cells and the FAN, particularly for tests C8 and C9.

**Figure 8.**
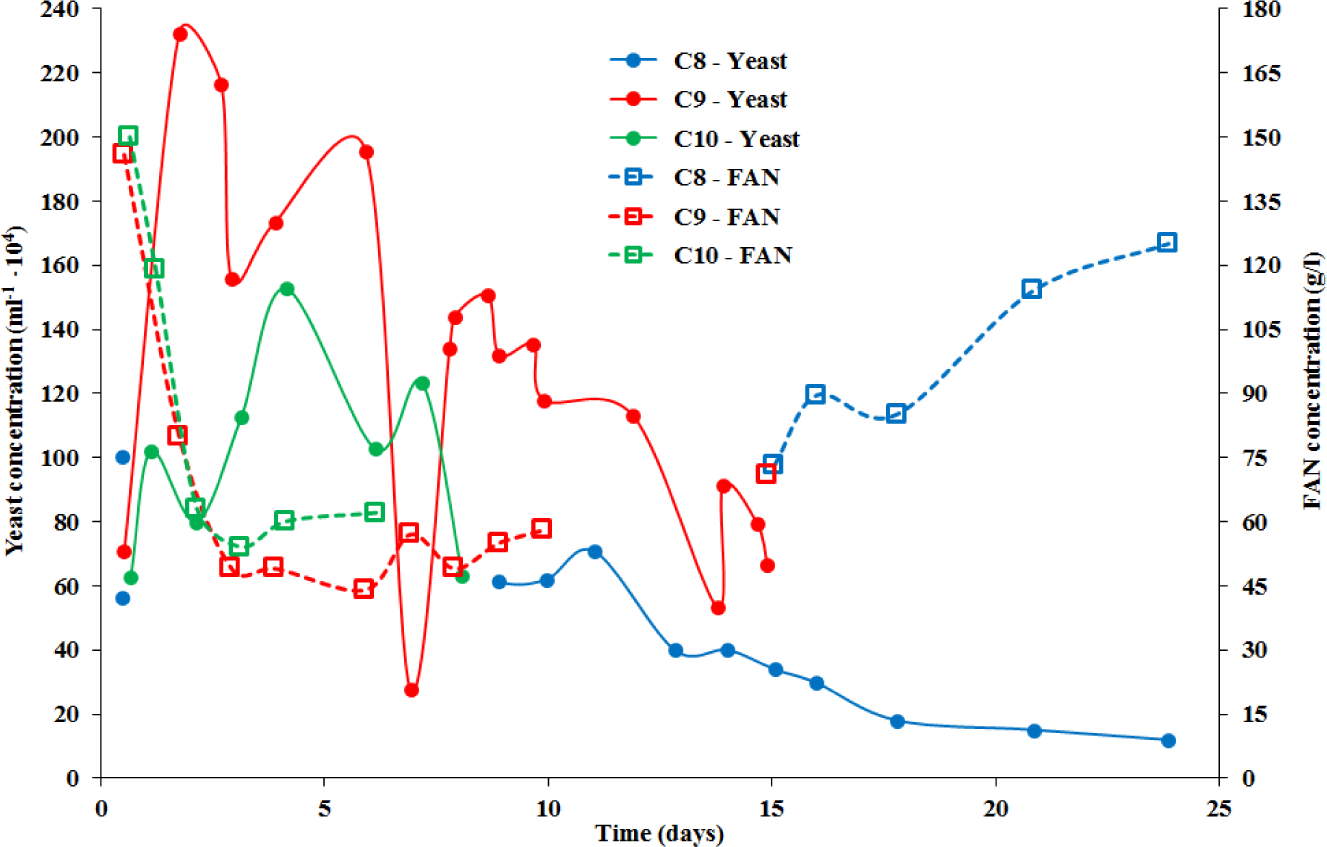
Tests C8, C9 and C10: concentration of yeast cells and FAN during fermentation.

Another interesting feature from Figure 8 is the apparently far greater concentration – on average, almost double – of alive yeast cells in test C9, despite very wide oscillations. Nevertheless, the FAN concentration curves from tests C8 and C9 link up towards the end of fermentation (precisely, the residence time in the open vessel) in test C9, *i.e.* about 15 days after yeast pitching. First, thus, yeast cells are partially inactivated by CHC processes in tests C8 and C10, which agrees with previous work by the authors (Albanese et al., 2015), with no apparent increase of lethality produced by the almost four-times longer treatment applied in test C10. Second, yeast cells in test C8, having been “activated” by the CHC process after their pitching, are more efficient in proline assimilation and irreversible degradation via the oxidase process, in comparison with test C9, so much that the simultaneously occurring effects of the reduction of alive yeast cells concentration and the activation of the survived ones compensate each other with respect to the impact on the FAN concentration. An hypothesis which agrees with the large difference in gluten concentration recovery during maturation shown in Figure 2 for tests C8 and C9, as well as with the observation that fermentation in test C8 started at least an hour earlier than in any other test.

Finally, it should be noted that the safe preservation of the beer wort in an open vessel well beyond the ordinary fermentation times, such as in test C8, is not just a trivial task, that is now achieved thanks to the CHC sterilization capabilities (Albanese et al., 2015, 2016). Similarly, it is remarkable that taste, aroma, flavor and foam stability of the finished beers from tests C8 and C10 do not appear to be adversely affected by the CHC treatment after yeast pitching, in agreement with what lately reported by Safonova and co-workers (Safonova et al., 2015).

In summary, what arises from tests C8, C9 and C10 is that the CHC treatment of the beer wort after yeast pitching for the purpose of gluten concentration reduction should be short enough (*e.g.* < 1 h) to avoid inactivation of a too large fraction of the yeast cells, while still generating much needed molecular oxygen. Furthermore, the CHC treatment will be coupled to longer than normal residence time in an open vessel (*e.g.* more than 20 days).

## 4. Conclusions

The new controlled hydrocavitation-assisted beer brewing technique (Albanese et al., 2016) provides another important advantage over conventional brewing technology, affording greatly reduced gluten concentration in the resulting beers. Eventually, in correspondence of suitable cavitation regimes identified in this study for barley malts, the amount of gluten found in beer is lower than the “gluten-free” threshold (20 mg/L). The relevance of this new route to gluten reduction arises from the fact that it allows to retain the same ingredients and recipes of standard beers, without the involvement of any chemical additives or proprietary techniques (such as filtration, ultrafiltration or enzymatic compounds), while preserving taste, flavor and aroma of the best craft beers. Moreover, all this comes without oxidation even after the most intense and prolonged cavitation processes as, within the range of CHC regimes used in the field of food applications, oxidation processes that are generally undesirable in liquid foods processing (Ngadi, Latheef, & Kassama, 2012) play a marginal role (Yusaf & Al-Juboori, 2014). Indeed, only the use of specific additives such as hydrogen peroxide allows achieving the needed extent of organics oxidation in applications such as water disinfection and remediation (Ciriminna, Albanese, Meneguzzo, & Pagliaro, 2016a).

Proline and its high levels in the hordein protein fraction of the barley grain (Deželak, Zarnkow, Becker, & Košir, 2014) is a key player for the gluten toxicity to coeliac patients or to the growing number of more mildly gluten-intolerant people (Benítez et al., 2016; Hager et al., 2014; Sollid, 2002; Uhde et al., 2016). Therefore, its possible assimilation, degradation and further reduction during fermentation and maturation will be very beneficial to the food safety of the finished beers.

Hydrodynamic cavitation was shown to be effective both during mashing and at the beginning of fermentation, *i.e.* after yeast pitching, with early operational guidance provided for both brewing stages. We ascribe this newly observed phenomenon to the formation of oxygen in the fermentation mixture becoming available to allow oxidase agents, such as enzymes (Schnitzenbaumer & Arendt, 2014) and yeasts (Procopio et al., 2013), to degrade proline. Supporting this insight, is the very recent finding that proline, whose concentration in the fermenting wort can be quite high, leading to the formation of fusel alcohols (Procopio, Krause, Hofmann, & Becker, 2013), shows an assimilation rate by yeast strains that increases in high-stress conditions due to the shortage of more easily assimilated amino acids, as well as with the increased availability of molecular oxygen, which is a scarce resource during anaerobic fermentation. Future developmental research will concern the measurement of proline, along with molecular oxygen at any relevant brewing stage, as well as the extension of the range of cavitation regimes, with identification of the operational parameters as function of brewing recipes. In the meanwhile, the first evidence of effective gluten reduction in some CHC-assisted beer brewing processes has been discovered and reported.

## Acknowledgements

We thank F. Martelli (CNR-IBIMET) and C. Capriolo (CNR-ICVBC) for continuous support in project management and technical help, respectively. L.A. and F.M. were partially funded by Tuscany regional Government under the project T.I.L.A. (Innovative Technology for Liquid Foods, Grant N°. 0001276 signed on April 30, 2014). The research was carried out under a cooperation between CNR-IBIMET and the company Bysea S.r.l. Joint international patent application No. PCT/IT/2016/000194, submitted on August 9, 2016, pending.

## Abbreviations

BIAB: Brew In the Bag
CN: Cavitation Number
FAN: Free Amino-Nitrogen
CHC: Controlled Hydrodynamic Cavitation
SG: Silica Gel
GRAS: Generally Recognized as Safe

